# The female intimate microbiome space

**DOI:** 10.1101/2025.02.26.640345

**Authors:** Leonore Vander Donck, Thies Gehrmann, Sarah Ahannach, Sarah Van den Bosch, Margo Hiel, Lize Delanghe, Camille Nina Allonsius, Eline Cauwenberghs, Irina Spacova, Eline Oerlemans, Stijn Wittouck, Ilke De Boeck, Gilbert Donders, Veronique Verhoeven, Sarah Lebeer

## Abstract

Despite its importance in health, the female skin microbiome remains understudied. In this study we explored microbial dispersal across intimate body sites, including the vagina, groins, breast and mouth. Microbial similarity correlated with physical proximity, suggesting dispersal influenced by hygiene or sexual activity. Notably, lactobacilli were unexpectedly abundant on breast skin. These findings highlight the need for research into microbiome dynamics and their implications for women’s health.

## Main section

The human skin is the largest epithelial surface for microbial interaction and a key barrier influenced by endogenous and exogenous factors^1–3^. Its complex topography and multiple layers contribute to a highly variable and phylogenetically diverse microbiome^4,5^. Most skin typical bacteria reside on the epidermis, but they are also present in deeper layers^2,6^. Different skin regions host distinct microbial communities: moist areas such as the groin favor *Staphylococcus* spp. and *Corynebacterium* spp., sebaceous regions such as forehead are rich in *Cutibacterium* spp., and dry areas such as forearm and elbow exhibit greater diversity, including *Corynebacterium* spp. and *Proteobacterium* spp.^3,4,7,8^.

Disruptions in the skin microbiota are linked to inflammatory skin conditions such as acne^9,10^, psoriasis^11^, and atopic dermatitis^9,12,13^. The microbiome plays a crucial role in skin barrier integrity^14^, modulating inflammatory responses^7^, wound healing, epidermal differentiation, and tight junction enhancement^2,3,15^. While lactobacilli are widely recognized as beneficial for gut and vaginal health, their role in skin health is only recently explored^14,16–19^. Though the skin’s nutrient-poor and oxygen-rich environment is unfavorable for lactobacilli, reduced levels have been associated with skin diseases in different observational studies^16,20–22^. As the skin is constantly exposed to external microbes, lactobacilli detected on the skin in observational studies may be transient members. They could for example be transferred from other body sites—particularly the vaginal microbiome, where they are abundant^23,24^. Lactobacilli could also be early colonizers of the skin upon vaginal birth^16,17^. However, studies linking vaginal and skin microbiomes of mother to children and within the same (female) individual are needed to confirm such dispersal patterns, but currently lacking.

The vaginal microbiome has gained research interest in relation to pregnancy, fertility, and infections, yet women remain underrepresented in scientific studies^25,26^. Gender biases in drug testing and clinical trials persist, leading to gaps in knowledge about the female skin microbiome. Although men and women have distinct skin characteristics—women generally have thinner skin, lower pH, and less sweat production—most skin microbiome studies do not distinguish between genders. Additionally, differences in hygiene practices contribute to a more diverse female skin microbiome^27–29^. Despite this, research rarely specifies whether samples come from men or women, instead summarizing data by body site.

The **Isala** citizen-science project addresses an important female microbiome knowledge gap by mainly investigating the vaginal microbiome. Initially, 3,345 women completed surveys and provided vaginal swabs via self-sampling^24^. In this study, we report on the second phase where women also donated samples of specific intimate skin sites. A subset of 293 participants, selected based on hormone intake, sexual activity, and contraceptive use, participated in this second phase. During the luteal phase of their menstrual cycle, they provided saliva (data already reported)^30^; a vaginal swab and skin swabs from the mouth area, breasts, and groin (reported here), as well as answered a large survey (Figure 1A). An overview of cohort selection and (intimate) hygiene characteristics can be found in supplementary figures 1 and 2. General cohort characteristics are described in Cauwenberghs *et. al*.^*30*^.

The bacterial DNA of the vaginal, groin, mouth and breast skin swabs and saliva samples was then analysed using the V4 region of the 16S rRNA gene to assess the microbiome of each individual’s intimacy related body site. In total 2,414 samples passed quality control, resulting in 51 million high-quality V4 amplicon paired reads. Similarly to the vaginal microbiome reads of the first phase of Isala ^24^, the reads were classified up to subgenus level with additional subgenera classification of the genus *Lactobacillus*^*24*^. An overview of the microbiome of each site for all participants can be found in supplementary figure 3. The vaginal microbiome was mainly dominated by *Lactobacillus crispatus, Lactobacillus iners* and *Gardnerella sp*. (also referred to as *Bifidobacterium vaginalis* in a newly suggested classification^31^), showing that in the vaginal microbiome, most participants (64%, 159/247 participants) are dominated by the same ASV in their vaginal microbiome as to their profile during the first phase of the Isala study^24^. All skin sites were mainly dominated by *Staphylococcus sp. a*nd *Corynebacterium sp*., which is consistent with the skin microbiome of moist skin sites. Interestingly, *Cutibacterium spp*. were detected in only eight skin samples, with a low average relative abundance (0,15%) given the sebaceous nature of the skin around the mouth and breasts^3,4^. However, this low abundance can be explained by the study design as it is known that the V4 region lacks in capturing *Cutibacterium spp*.^32,33^. V4 was still selected here as target variable regions to be able to compare skin sites with vaginal samples (especially *Lactobacillus* taxa) and because *Corynebacterium* was not the target bacterium of this study. The salivary microbiome of this cohort was a conserved community of 12 genera, mainly consisting of *Streptococcus, Veillonella* and *Prevotella*, as previously reported by Cauwenberghs *et al*.^*30*^.

We then mapped the community structure of all body sites in a two-dimensional space using t-distributed stochastic neighbour embedding (t-SNE) at ASV level Figure 2B)^34^. Samples clustered on site (figure 2B), which is in accordance with previous findings^2,35^. When plotting the relative abundance of the top (sub)genus on ASV level, we mainly observed a high relative abundance of a singly top (sub)genus in the vaginal microbiome, while the salivary microbiome had an overall lower average relative abundance (figure 1E), revealing a core salivary microbiome^30^. To uncover insights into the dispersal of microbiota through different body sites of one participant, the different samples of two participants were visualized throughout the microbiome space as examples (Figure 1B, 1C, 1D). The samples of a single participant were clearly placed on a trajectory through the microbiome space showing the dispersal of more site-specific taxa to nearby body sites. For example, the vaginal microbiome of participant 1 was almost singularly dominated by *L. crispatus*, which resulted in the immense dispersal of *L. crispatus* to the groin skin, with a relative abundance of more than 70%. This almost resembled a more typical vaginal microbiome, with only little *Staphylococcus* and *Corynebacterium* present. The groin skin of participant 2 showed less resemblance to her vaginal microbiome, which was dominated by *L. iners*, but still contained more than 20% of both *L. iners* and *L. jensenii* which are found in her vaginal microbiome. The microbiome of the skin of the mouth of participant 2 was also more similar to the skin microbiome of the breasts, while the mouth skin of participant 1 contained also more typical salivary bacteria such as *Veillonella* and *Streptococcus*. These data suggest more dispersal of vaginal fluid to the groins and the saliva to the skin of the mouth in participant 1 compared to participant 2, although it remains to be explored whether these found on the skin sites are the same strains that are found in the vagina or saliva, to confirm certain dispersal.

To further substantiate possible dispersal to nearby sites, the distance between the microbiome of each site was determined (Figure 1H). The distance between microbiomes correlated linearly with physical distance, following a ‘vertical flow’ across body sites; from the vagina to the groin skin, then to the breast skin, the skin around the mouth, and finally, the salivary microbiome. This was visible in the number of taxa on subgenus level overlapping between sites. The groin skin was an overlap between the vaginal microbiome and skin typical microbiome taxa, while the skin around the mouth showed an overlap between the salivary microbiome and the typical skin microbiome taxa. Unexpectedly, the skin around the breasts contained a notable amount of typical vaginal lactobacilli, with an average relative abundance of 3% on the breasts, compared to an average abundance of 1.9% on the skin around the mouth. We hypothesize a possible distribution of vaginal lactobacilli to other body sites influenced by sexual activity and (intimate) hygiene practices, although the lactobacilli could also be endogenously enriched for example due to increased hormone levels and oestrogen receptors in breasts^36^, sustaining the presence of lactobacilli similarly as in the vaginal niche. While no significant associations between hygiene habits and recent sexual activity and the skin microbiome sites in our cohort were found (Supplementary Figures 4 and 5), the elevated abundance of lactobacilli on intimate sites is in accordance with Howard B. *et al*. who found more *Lactobacillus* sp. on the buttocks of women than on the face and forearm^37^. Lactobacilli were also already found on the female hands, with a higher abundance on the dominant hand, hinting towards dispersal of the vaginal microbiome^38^. A possible dispersal due to sexual activity has also been observed in cohabiting couples, where *Lactobacillaceae* were found on the inner thighs of male partners^39^. This provides new biological insights in bacterial dispersal, but could also be relevant in sexual assault cases, where the microbiome has recently gained interest as biological evidence^40–42^.

The dispersal of bacteria across different sites on the human body is an understudied phenomenon. Here, we demonstrate the relationship between microbiomes at different physical proximities within the same individual. Further studies with higher taxonomic resolutions are needed to elucidate the actions that effect the dispersal of site specific microorganisms and the impact on the affected body sites. Specifically, the influence of the distribution of lactobacilli in relation to sexual habits or oestrogen levels is of interest due to its probiotic nature.

## Materials & Methods

### Study cohort and data collection

The study was approved by the Ethical Committee of the Antwerp University Hospital/University of Antwerp (B300201942076) and registered online at ClinicalTrials.gov with the unique identifier NCT04319536. 293 participants were selected from the larger cohort of 3,345 women based on hormone intake (no hormones, IUD, combination pill), condom usage (yes, no), frequency of sexual contact (very frequent, frequent, saldo, never) and number of partners (multiple, one, none). Participants were contacted again and agreed to participate to continue in a second phase where over the course of two menstrual cycles samples were collected. Participants collected vaginal swabs at three timepoints in one menstrual cycle. During the luteal phase of the first menstrual cycle, they also collected a saliva sample and swabbed the skin around the mouth, around the breasts and of the groin. 258 participants collected all skin samples, 247 participants provided saliva samples and 246 participants provided all six vaginal swabs.

### Sample collection saliva and skin swabs

Participants provided saliva samples by spitting into a sterile plastic cup and twisting the eNAT® swab in the saliva. For the sampling of the skin of breast (n=231) eNAT® swabs (Copan, Brescia, Italy) were used for microbiome profiling. Participants moistened the swabs with a sterile pre-moisture buffer (50 mM Tris buffer [pH 7,6], 1 mM EDTA [pH 8,0], and 0,5% Tween-20) and gently rubbed the area for 30 seconds to collect sufficient biomass. The skin of mouth (n=251) and groin (n=251) the participants moisturized the eNAT® swab with 10x PBS and rubbed it for 30 seconds on a 20 cm^2^ area.

### Sample collection vaginal swabs

The vaginal swabs were collected and processed as mentioned in Lebeer S. *et. al*. (2023)^24^.

### 16S rRNA amplicon sequencing, reference database and quality control

All samples were processed and analysed as described in Lebeer S. *et. al*. (2023)^24^.

### Statistical analyses

Statistical analyses used to determine the associations between microbial community composition and survey data were performed as described in Lebeer S. *et al*. (2023)^24^.

## Data availability

The sequencint data is available at the European Nuceotide Archive under bioproject PRJEB86101.

## Author contributions

S.A., G.D., V.V., E.O, S.W and S.L. designed the study and worked on the conceptualization of the research project. S.A. and S.L. worked on the survey set-up and L.V.D., T.G., S.A., I.S. cleaned the answers. S.A., L.V.D., E.C. and L.D. carried out the experimental and logistical work. I.D.B and E.C were responsible for the biobanking of all collected samples. T.G. and M.H. processed the sequencing data and performed the biostatistical analyses. L.V.D., T.G., S.V.B., M.H., S.A. and S.L. worked on the visualizations. L.V.D., T.G., S.A., S.V.D., M.H., C.N.A, and S.L. contributed to the interpretation of the results. L.V.D., S.A., T.G. and S.L. were responsible for the science communication to the participants in layman’s terms. L.V.D., S.A., T.G., S.V.B., M.H., S.L. wrote the original manuscript. All authors contributed to reviewing and editing of the final manuscript

---

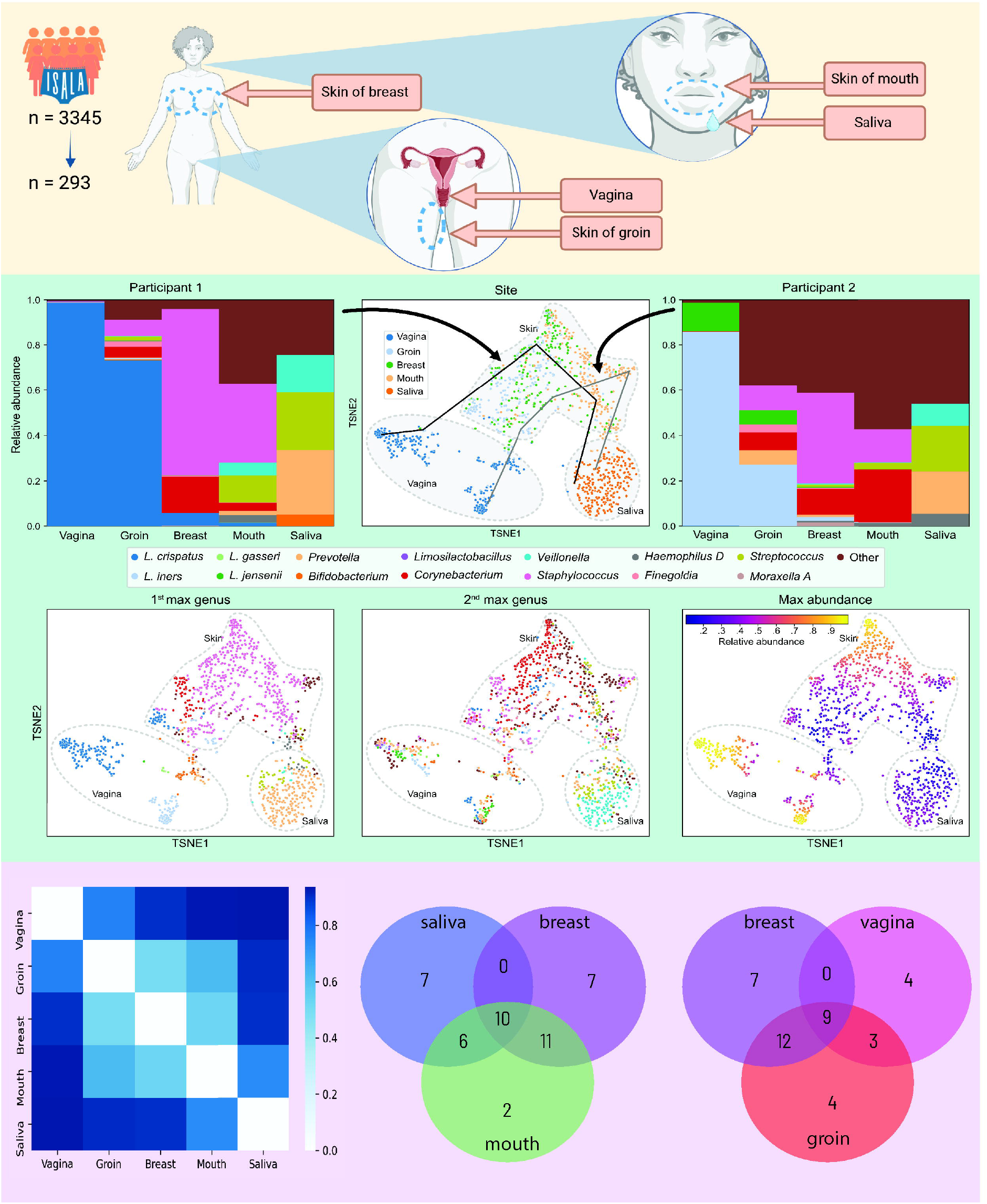

