## Supplementary tables and figures for "The female intimate microbiome space"

Table 1: Distances between the microbiomes of participant 1

| site<br>site | vaginal | skin_groin | skin_breast | skin_mouth | saliva |
|---|---|---|---|---|---|
| **vaginal** | 0.000000 | 0.995511 | 0.995171 | 0.995604 | 0.987699 |
| **skin_groin** | 0.995511 | 0.000000 | 0.348779 | 0.291753 | 0.943697 |
| **skin_breast** | 0.995171 | 0.348779 | 0.000000 | 0.236848 | 0.947357 |
| **skin_mouth** | 0.995604 | 0.291753 | 0.236848 | 0.000000 | 0.940363 |
| **saliva** | 0.987699 | 0.943697 | 0.947357 | 0.940363 | 0.000000 |

Table 2: Distances between the microbiomes of participant 2

| site<br>site | vaginal | skin_groin | skin_breast | skin_mouth | saliva |
|---|---|---|---|---|---|
| **vaginal** | 0.000000 | 0.992755 | 0.991763 | 0.990588 | 0.994670 |
| **skin_groin** | 0.992755 | 0.000000 | 0.338816 | 0.227453 | 0.990202 |
| **skin_breast** | 0.991763 | 0.338816 | 0.000000 | 0.359915 | 0.990039 |
| **skin_mouth** | 0.990588 | 0.227453 | 0.359915 | 0.000000 | 0.996262 |
| **saliva** | 0.994670 | 0.990202 | 0.990039 | 0.996262 | 0.000000 |

### Supplementary Figures

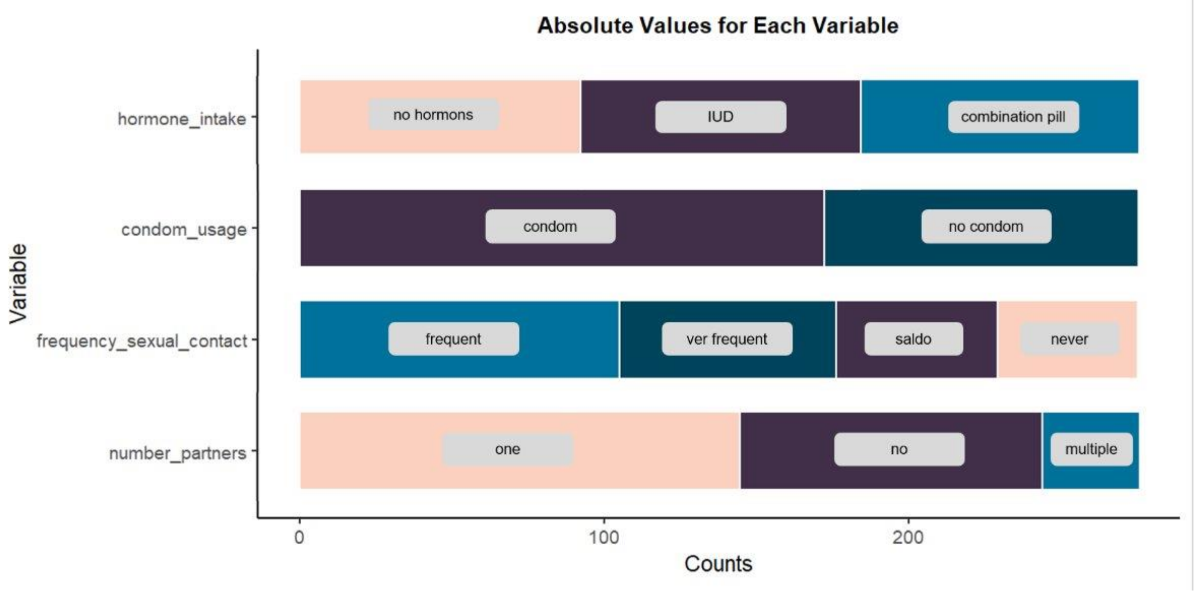


Figure 1: Selection of flow 2 cohort within the flow 1 Isala cohort

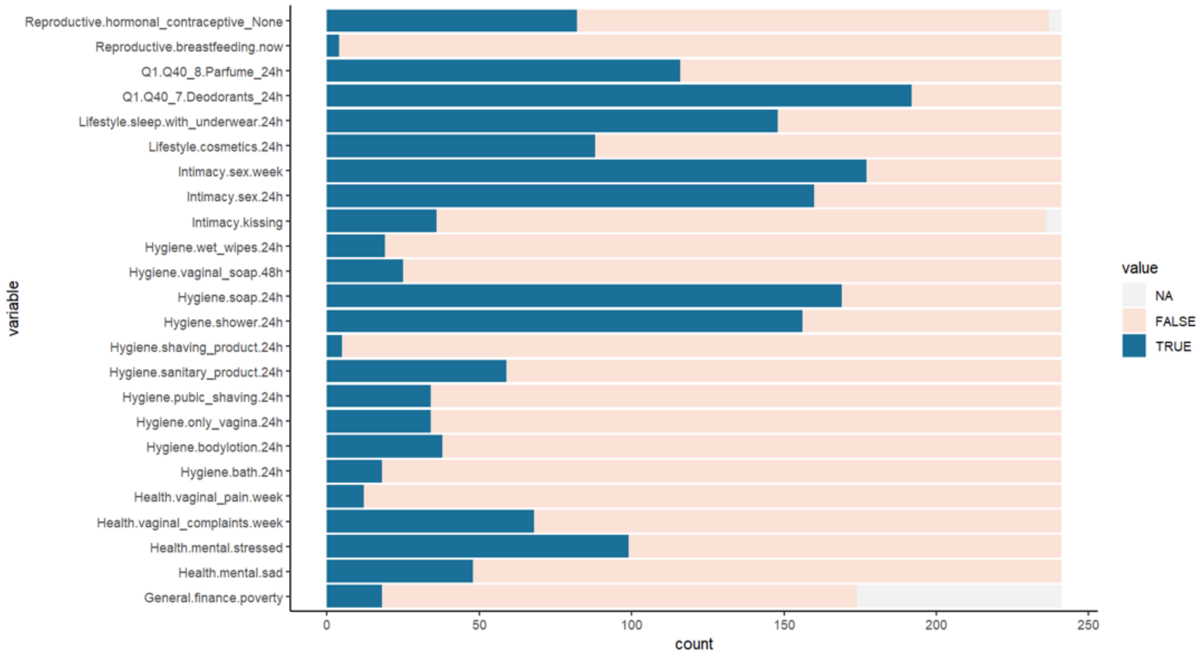


Figure 2: Cohort characteristics of the (intimate) hygiene habits

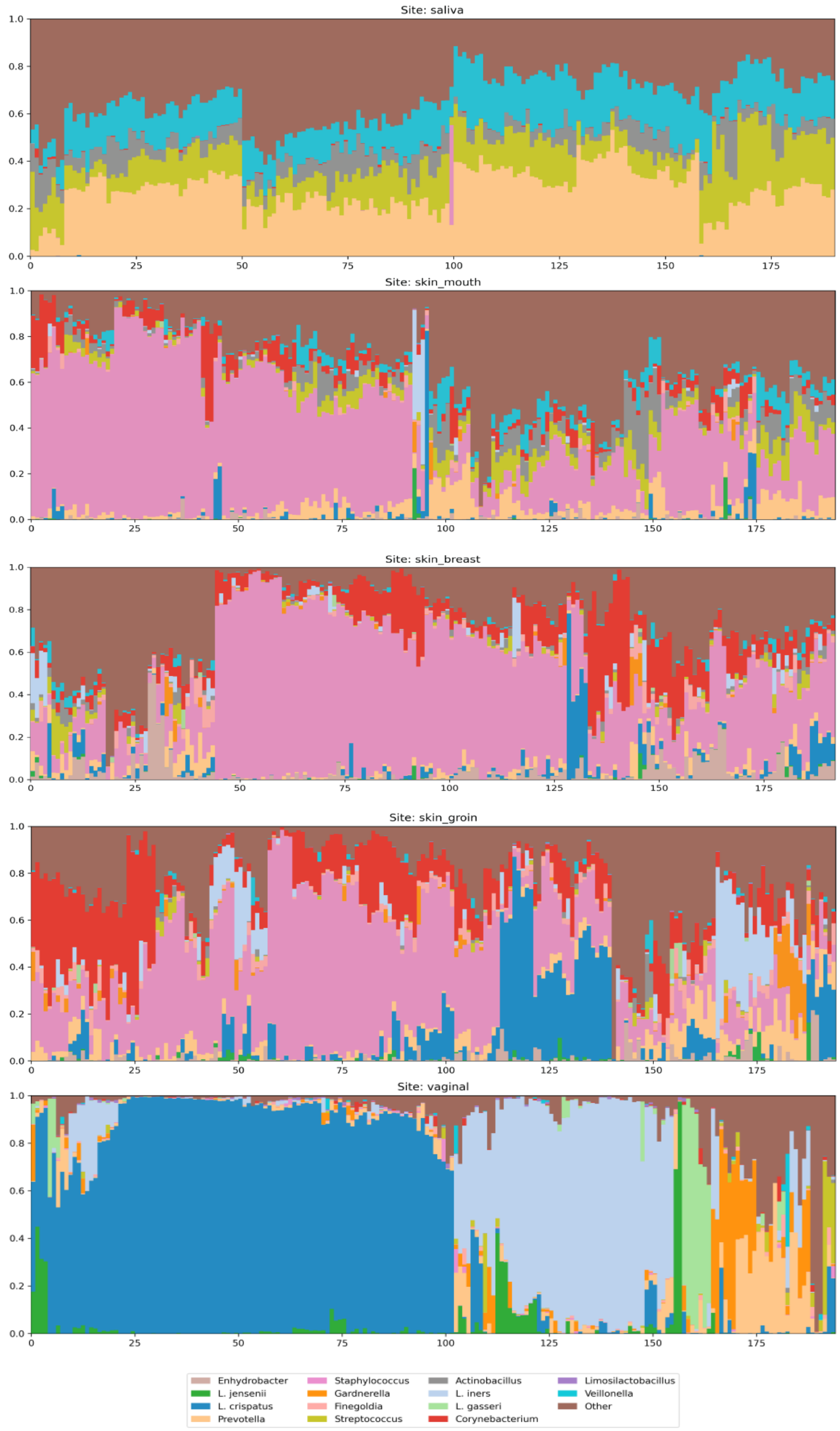
Site: saliva
Site: skin_mouth
Site: skin_breast
Site: skin_groin
Site: vaginal
Enhydrobacter
L. jensenii
L. crispatus
Prevotella
Staphylococcus
Gardnerella
Finegoldia
Streptococcus
Actinobacillus
L. iners
L. gasseri
Corynebacterium
Limosilactobacillus
Veillonella
Other

24

Figure 3: Barplots showing the overview of the saliva, skin of the mouth, skin around the breasts, groin skin and vaginal microbiome of the cohort.

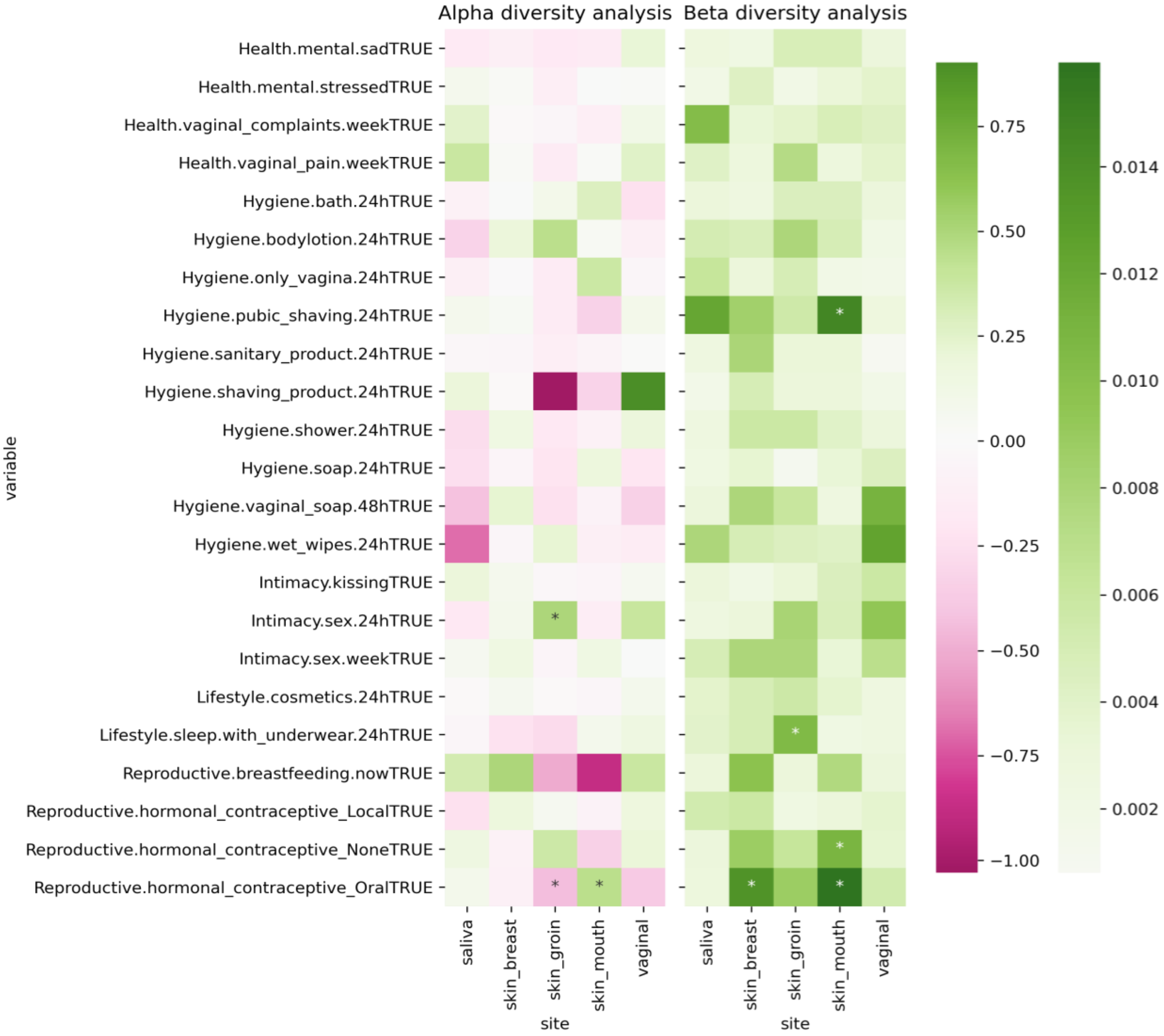


Figure 4: Associations of (intimate) hygiene characteristics of the cohort on different levels with the salivary, skin and vaginal microbiome: The effect on the alpha diversity (Shannon index) of the samples on different sites and the effect on beta diversity between the samples (Adonis test) on different sites. * represent significant associations ($q < 0.05$). For the individual taxa tested, the number refers to the number of pipelines that indicate a significant association: ALDeX2, ANCOM-BC, DESeq, limma, Maaslin2 and a linear regression on the CLR-transformed abundance data. All tests were adjusted for multiple comparisons with Benjamini–Hochberg procedure.

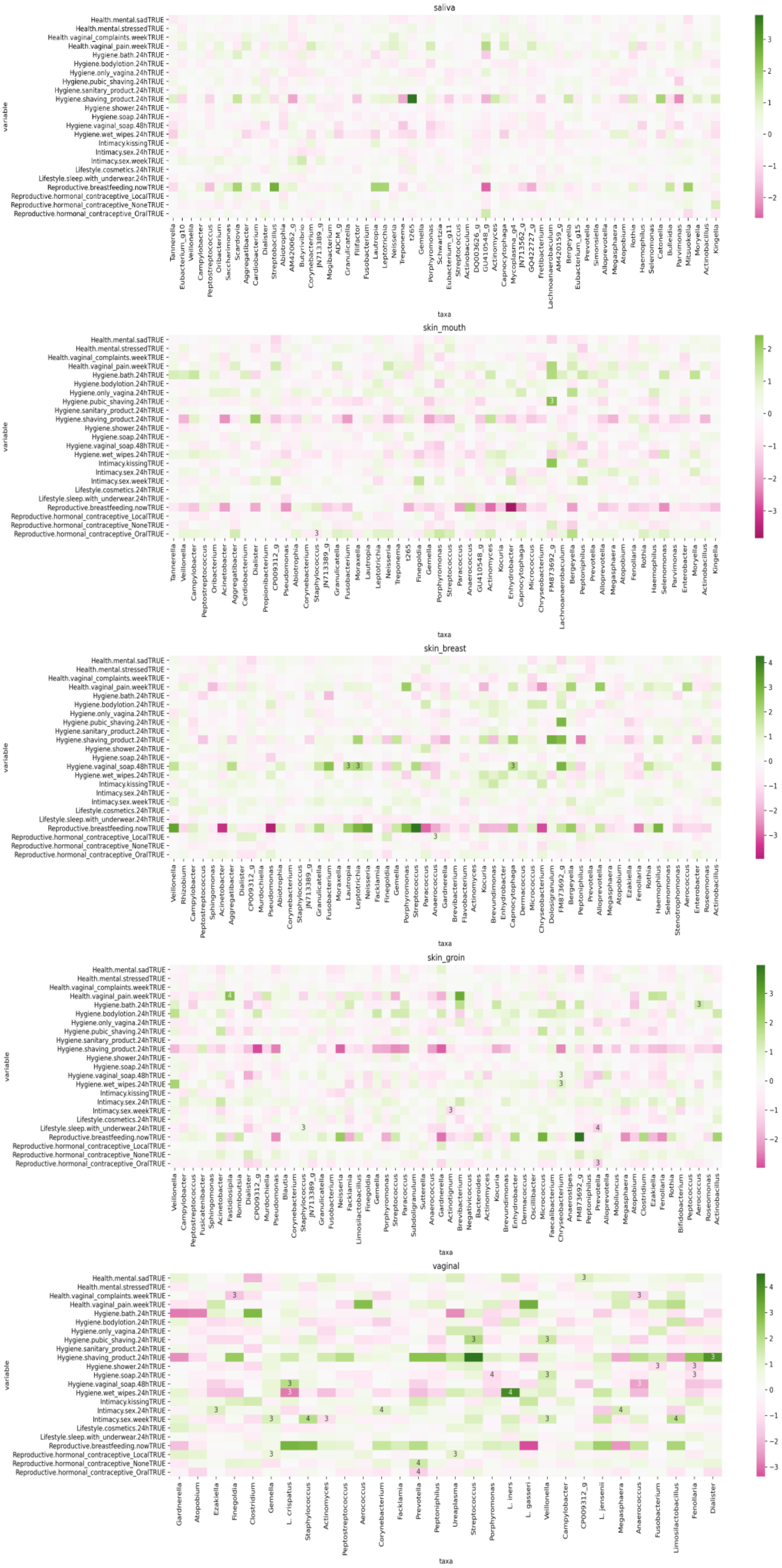
saliva
skin_mouth
skin_breast
skin_groin
vaginal
variable
taxa
Health.mental.sadTRUE
Health.mental.stressedTRUE
Health.vaginal_complaints.weekTRUE
Health.vaginal_pain.weekTRUE
Hygiene.bath.24hTRUE
Hygiene.bodylotion.24hTRUE
Hygiene.only_vagina.24hTRUE
Hygiene.pubic_shaving.24hTRUE
Hygiene.sanitary_product.24hTRUE
Hygiene.shaving_product.24hTRUE
Hygiene.shower.24hTRUE
Hygiene.soap.24hTRUE
Hygiene.vaginal_soap.48hTRUE
Hygiene.wet_wipes.24hTRUE
Intimacy.kissingTRUE
Intimacy.sex.24hTRUE
Intimacy.sex.weekTRUE
Lifestyle.cosmetics.24hTRUE
Lifestyle.sleep.with_underwear.24hTRUE
Reproductive.breastfeeding.nowTRUE
Reproductive.hormonal_contraceptive_LocalTRUE
Reproductive.hormonal_contraceptive_NoneTRUE
Reproductive.hormonal_contraceptive_OralTRUE

Figure 5. Associations of (intimate) hygiene characteristics of the cohort on the level of eigentaxa within the salivary, skin and vaginal microbiome.
